# A continuum of CAM phenotypes in the carnivorous plant genus *Pinguicula* (Lentibulariaceae)

**DOI:** 10.64898/2026.07.22.740078

**Authors:** Daniel W.L. Mok, Robert VanBuren, Kadeem J. Gilbert

## Abstract

**Premise:** *Pinguicula* are carnivorous plants occupying a wide range of habitats, from wetlands to barren rock cliffs, where CAM photosynthesis was recently discovered in several species. This ecological and physiological diversity makes the genus a promising system for exploring the environmental drivers and evolutionary transitions underlying variation in CAM. Here, we present a physiological survey of CAM with a particular focus on the Mexican Clade, which contains the recently identified CAM species.

**Methods:** We conducted a carbon isotope survey of live and herbarium specimens to identify candidate CAM species. We then measured diurnal gas-exchange patterns and changes in leaf titratable acidity to quantify CAM activity.

**Results:** *Pinguicula* species exhibited a variety of photosynthetic phenotypes. *Pinguicula vulgaris* showed no detectable evidence of CAM, whereas *P. cyclosecta* and *P. moranensis* demonstrated a C_3_-CAM physiology with most net assimilation via C_3_. *Pinguicula martinezii* also showed the same C_3_-CAM physiology under well-watered conditions but had strong facultative induction of CAM with drought. *Pinguicula agnata* displayed net assimilation via CAM. Titratable acidity had a positive correlation with δ^13^C within the species we sampled.

**Conclusions:** Our results support the existence of a CAM physiological continuum within *Pinguicula*. Notably, closely related species encompass physiologies that span the large variation of CAM. This diversity provides a rare opportunity for future studies to investigate the evolution of CAM in closely related species, including differences and similarities between differing states across the continuum.

## INTRODUCTION

*Pinguicula* (Lentibulariaceae), also known as butterworts, are a genus of carnivorous plants composed of small rosettes with diverse leaf morphology and coloration (Figure 1). They have sticky-trap carnivorous leaves covered in sessile and stalked secretory glands (Lampard et al., 2016; Roccia et al., 2016). The genus is native to all continents except Australia and Antarctica, with the temperate Northern Hemisphere proposed as its geographic origin (Domínguez et al., 2024). The highest diversity of species is found in Mexico, which harbors nearly half the genus. Given this broad geographic distribution, *Pinguicula* are found in a diverse array of habitats. Temperate *Pinguicula* species grow in areas with abundant year-round water, like bogs or wet, rocky seeps (Legendre, 2000; Heslop-Harrison, 2004; Shimai, 2017; Shimai et al., 2021). This distribution aligns with the well-supported habitat distribution for carnivorous plants predicted by the cost-benefit model of plant carnivory, which predicts that carnivorous plants should be restricted to areas with low nutrient availability and abundant water and light (Givnish et al., 1984; Ellison and Gotelli, 2001; Givnish, 2015). In such habitats, the benefits of carnivory to photosynthesis (e.g., increased nitrogen) outweigh the physiological costs associated with being carnivorous (Givnish et al., 1984; Ellison, 2006; Pavlovič and Saganová, 2015).

**Figure 1:**
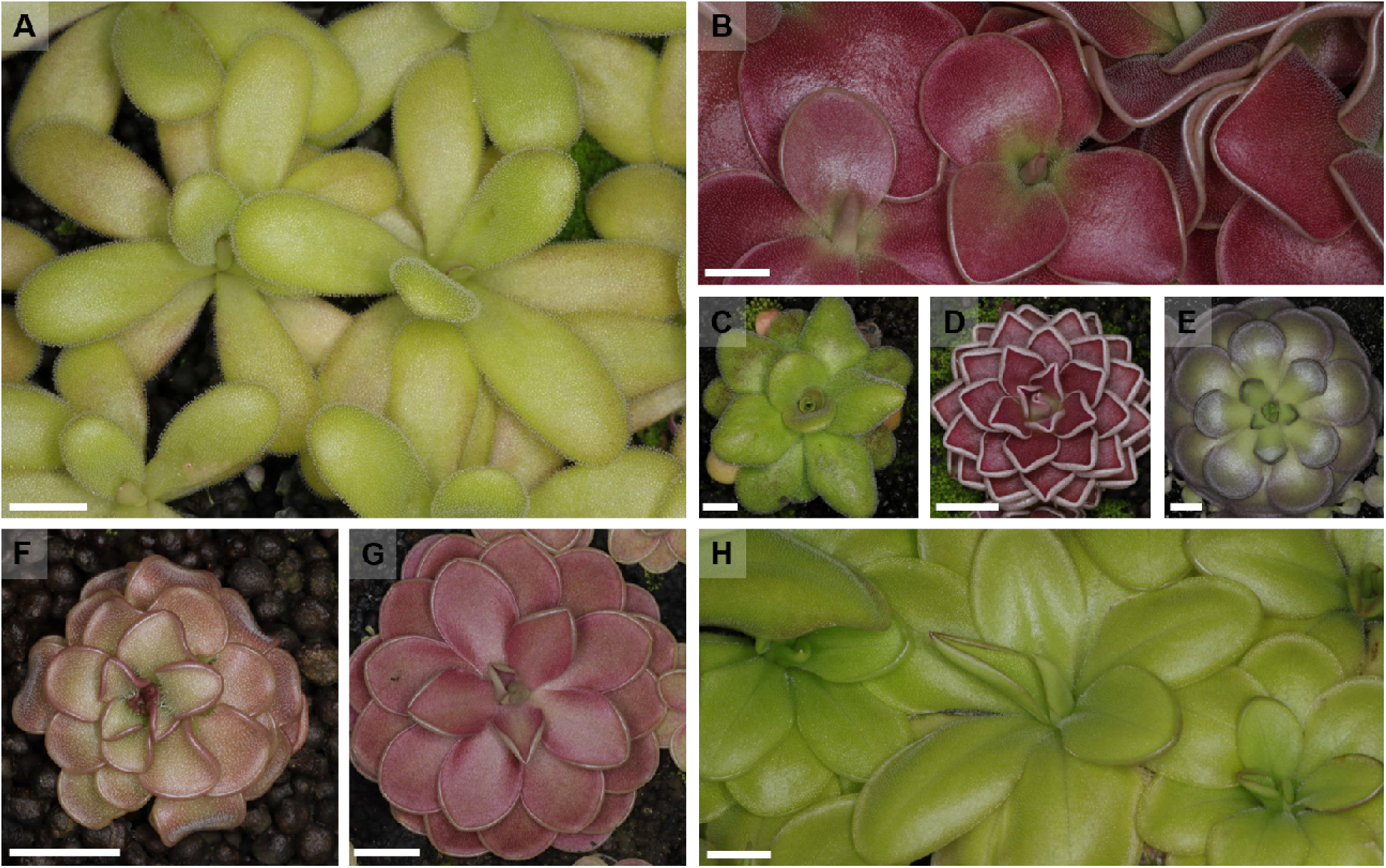
Images showing a variety of Mexican *Pinguicula* species growing in our living collections: A - *P. agnata*, B - *P. moranenesis*, C - *P. martinezii*, D - *P. ehlersiae*, E - *P. cyclosecta*, F - *P. rotundiflora*, G - *P. jaumavensis*, H - *P. zecheri*; White bar = 1 cm

However, in Mexico, many *Pinguicula* species grow on trees or bare, near-vertical rock cliffs, where water availability is often limited and seasonal drought can impose substantial water stress (Mata-Rosas et al., 2020; Rueda-Almazán et al., 2021). Such habitats are inconsistent with the distribution predicted by cost-benefit models. If plant carnivory requires abundant water, how have these xeric-dwelling species maintained carnivory while also adapting to low-water environments?

Crassulacean acid metabolism (CAM) has been proposed as a strategy for overcoming water limitations in Mexican *Pinguicula* (Miloslav Studnička, 1991; Pavlovič, 2022) and has recently been demonstrated in the genus (Fleck et al., 2026). CAM shifts carbon assimilation to the night, storing it as malic acid (Osmond, 1978; Okita, 1996; Lüttge, 2004). Stomata may close during the day, and the acid is converted back to CO_2_, minimizing water loss under the higher transpirational pressures (Osmond, 1978; Cushman, 2001; Lüttge, 2004). This makes CAM potentially up to six times more water-use-efficient relative to C_3_ photosynthesis (Winter et al., 2005; Yang et al., 2015). Because water limitation is a common selective pressure for plants, CAM provides an effective physiological solution (Borland et al., 2011; Heyduk, 2022) and has consequently evolved over 60 times independently in vascular plants (Gilman et al., 2023).

CAM, however, is distinct in its phenotypic variation, sometimes termed the ‘CAM continuum’, which orders species based on the relative contributions of CAM towards the plant’s overall carbon assimilation (Silvera et al., 2010; Edwards, 2019). Species vary in both the extent to which they use CAM, either facultatively (e.g., inducible under stress) or constitutively, as well as the proportion of lifetime carbon gain derived through CAM (Smith and Winter, 1996; Winter, 2019; Yuan et al., 2020).

The community’s investigation of how this continuum evolved has been limited by two major hindrances: (1) Most CAM-evolving lineages are extremely species-rich (i.e., Orchidaceae (Silvera et al., 2009; Givnish et al., 2015), Cactaceae (Arakaki et al., 2011), Euphorbiaceae (Horn et al., 2012), Bromeliaceae (Crayn et al., 2004; Silvestro et al., 2014)), and thus thorough sampling of taxa is difficult; and (2) Many CAM-evolving lineages also lack relevant phenotypes along the CAM continuum, making it difficult to reconstruct evolutionary trajectories (Heyduk et al., 2019; Edwards, 2023). Since evidence for both non-CAM (C_3_) and facultative CAM states have previously been demonstrated in *Pinguicula* (Fleck et al., 2026), identifying additional species with varying CAM physiology could make *Pinguicula* a highly informative lineage for studying CAM evolution.

A combination of stable isotopes, acid titrations, and gas exchange measurements are used to determine CAM phenotypes of plants with unknown physiology. Carbon isotope (δ^13^C) surveys of leaf material can distinguish C_4_ and CAM plants from C_3_ species (Osmond et al., 1973; Winter and Holtum, 2002; Sage et al., 2011; Gilman et al., 2023) since PEPc, the primary carbon fixing enzyme in CAM and C_4_, discriminates less against ^13^CO_2_, leading to tissues with less negative δ^13^C (Farquhar and Lloyd, 1993). For CAM, δ^13^C less negative than -20‰ is an indicator for a majority of total CO_2_ assimilation occurring via CAM at night (Winter and Holtum, 2002). However, plants with low amounts of CAM expression can be masked in these surveys as their δ^13^C are often nested within the typical δ^13^C of C_3_ plants (Messerschmid et al., 2021) and thus, require physiological validation. Since CAM plants accumulate malic acid at night and decarboxylate the acid to release CO_2_ during the day (Osmond, 1978; Okita, 1996), differences in leaf tissue titratable acidity between dawn and dusk can also identify CAM (Winter and Smith, 2022). Lastly, measurements of gas exchange reliably identify the relative contribution of CAM towards a plant’s total carbon assimilation. CAM plants have distinct diurnal gas exchange patterns that readily distinguish them from C_3_ or C_4_ plants (Osmond, 1978; Winter, 2019). Together, the combination of these three techniques allows for the accurate identification and description of phenotypes spanning the CAM continuum.

Prior physiological work in *Pinguicula* investigated the induction of CAM (Fleck et al., 2026), but the broader distribution of CAM phenotypes across the genus, including the potential for constitutive CAM under well-watered conditions, remains uncharacterized. Here, we show that CAM expression varies across *Pinguicula*, specifically in the Mexican Clade, including previously unrecognized CAM phenotypes in the group. We first conducted a broad δ^13^C survey of herbarium and living specimens to identify candidate CAM species, then used continuous gas exchange measurements and titratable acidity assays to better characterize CAM physiology in representative taxa spanning the isotopic range we observed. We characterized multiple candidate CAM species in the genus and several species with varying degrees of CAM intermediacy, showing in particular, the existence of a CAM continuum within the genus *Pinguicula*. This physiological diversity highlights the potential for *Pinguicula* as an emerging model system for understanding CAM evolution.

## MATERIALS AND METHODS

### δ^13^C surveys

To create a broad survey of potential CAM species across *Pinguicula*, herbarium specimens were sampled for δ^13^C from the University of Michigan Herbarium (51, from MICH), or donated from the Billie L. Turner Plant Resources Center (14, from TEX), Lundell Herbarium (3, from LL), Missouri Botanical Gardens (6, from MO), Herbarium of the Arnold Arboretum (1, from A), and Harvard University Herbaria (2, from GH). We also compiled δ^13^C from prior studies where values for *Pinguicula* were reported: 20 from Klink et al. (2019); 6 from Lüttge, (1983); and 2 from Osmond et al., (1975). We additionally sampled leaf tissue from select species with sufficient material in our living collections at Michigan State University. In total, our δ^13^C survey used 77 herbarium specimens, 28 previously reported values, and 36 cultivated specimens, representing a total of 56 species across the *Pinguicula* phylogeny, with an emphasis on the Mexican Clade. Leaf tissue from herbarium and live specimens (dried at 60°C for 48 h) were weighed and encapsulated in 3.5 mm x 5 mm tin capsules. For each sample, we used between 0.3–0.7 mg of dried tissue for our δ^13^C assays, which were conducted by the Washington State University stable isotope facility (https://labs.wsu.edu/isotopecore/).

### Plant growing conditions

The plants in our living collections were purchased from specialty carnivorous plant stores or obtained from private growers. Mexican *Pinguicula* species were grown in 30 cm x 60 cm trays with plants spaced approximately 2 cm apart in a ∼15 mm depth of Fluval Stratum (Fluval, Baie-D’Urfe, QC, Canada, https://fluvalaquatics.com/), with a 12-h day/night period with photosynthetic photon flux densities (PPFD) of ∼250–300 μmol photons m^−2^ s^−1^ at the leaf level, 26°C day/22°C night temperatures, and 70% relative humidity. We grew the Mexican and temperate species with different substrates and watering regimes to better match the environmental conditions of their respective native habitats. Our temperate species, *P. vulgaris,* was grown in 2.5 inch pots with Suregro Mix (Sure Grow AG Products, Comanche, Texas, https://suregrowag.com/), in a tray under the same environmental conditions. To maintain our Mexican species under well-watered conditions, we gave each tray ∼450 mL of distilled water three times a week. *Pinguicula vulgaris* pots were kept well-watered by maintaining standing water in the trays. Plants were fertilized every two weeks with the same volume of a 50:50 distilled water: nutrient solution mix.

### Gas exchange measurements

Diurnal gas exchange measurements to measure CAM activity were taken using a LI-6800 system with the small plant chamber head attachment (LI-COR Biosciences, Lincoln, Nebraska, USA; www.licor.com). We selected species for gas exchange measurements based on availability in our living collection, being an adequate size to provide enough photosynthetically active tissue and having δ^13^C values that spanned the range of our survey results. Our specimens were not large enough to have individual leaves clamped in a traditional clamp-on-cuvette and the leaves were also too fragile, hence the use of the small plant chamber attachment. We measured *P. vulgaris*, from the North Temperate Zone, as a hypothesized temperate C_3_ species, *P. cyclosecta* and *P. moranensis*, from Mexico, as species with δ^13^C values in the C_3_ range, and *P. agnata*, from Mexico, a species with CAM-like δ^13^C values. We also took gas exchange measurements over a 20-day drought time course to test for facultative CAM in the Mexican species *P. martinezii*. For each species, we measured gas exchange on one individual.

Carnivorous plants tend to have overall low photosynthetic rates relative to non-carnivorous plants (Hájek and Adamec, 2010; Ellison and Adamec, 2011; Adamec et al., 2021) and the small plant chamber allowed us to maximize the amount of photosynthetic tissue in the chamber for more accurate measurements under low assimilation rates. To overcome issues of evaporation and respiration from substrate, we wrapped the surface of the container tightly around the base of the plant with several layers of parafilm and a silicone putty ring to reduce the area of the pot exposed to the chamber. This technique allowed us to more accurately capture the diurnal CO_2_ assimilation patterns of our species despite low photosynthetic rates. Starting in the dark period (18:00 h), continuous measurements were taken every five minutes with ambient CO_2_ concentrations and followed the temperature cycling and lights of the growth chamber.

Plants placed into the chamber were considered well-watered on the first day, and subsequent days were considered as days of water-withholding. Plants used for gas exchange were imaged and we used ImageJ (Schneider et al., 2012) to estimate a projected two-dimensional leaf surface area for the area-based gas exchange estimates presented. Though this could lead to overestimation of CO_2_ assimilation rates for non-flat and overlapping leaves, our focus in this study is on the diurnal trajectory of gas exchange rather than the absolute rates.

In order to visualize the measured trajectories, we plotted net CO_2_ assimilation rate (*A*), over time. To quantitatively assess the patterns seen in our gas exchange data, we integrated the area under the curve to determine the proportion of CO_2_ contributed via CAM. For species which had entirely net positive *A* throughout the dark period, we integrated *A* over paired 12 h dark and light periods to calculate the proportional net contribution of each to total daily carbon gain. For species without net positive dark period *A*, we estimated a hypothetical ‘no-CAM’ respiratory baseline for each night by fitting a linear model of *A* against time using anchor points from the beginning and end of the dark period. We then integrated both observed and estimated respiration to calculate the proportion of respiratory CO_2_ recycled via CAM.

### Titratable Acidity Measurements

In a separate experiment, we measured the tissue acidity of 20 representative *Pinguicula* species by sampling fresh leaf tissue from well-watered plants at dawn (06:00 h) and dusk (18:00 h). At each time point we sampled three clonal individuals per species. Tissue was immediately frozen in LN_2_ and stored at -80°C until processing. We incubated ∼25–90 mg of fresh leaf tissue in 1.4 mL of 20% EtOH (V_total_) in a water bath at 65°C for 2 hours and used 20% EtOH samples as a negative control. After incubation, all samples were briefly vortexed, and a 1 mL aliquot (V_extract_) was transferred to a new 1.5 mL tube containing 50 L of 0.004% solution of the pH indicator Bromothymol Blue. Solutions were titrated in 1 to 5 L increments of 0.005 M NaOH, until pH 7 indicated by the reaction solution turning a blue-ish green (μL _sample_). From the control EtOH tube, 1 mL was also titrated to pH 7 and the resulting volume (μL _control_) was subtracted from the samples. Titratable acidity on a fresh weight basis was calculated according to the following equation.

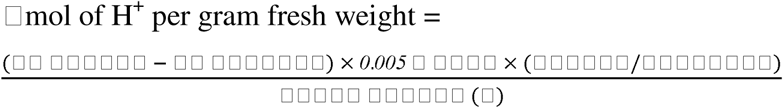

We mapped titratable acidity measurements onto the most recent phylogeny for the genus (Liu et al., 2026) with additional taxa (Shimai et al., 2021) to assess whether diel changes in acidity were associated with phylogenetic relationships. To calculate the maximum difference in acidity (ΔH^+^), we subtracted titratable acidity values at 18:00 h from 06:00 h. For each species independently, a Welch’s t-test was used to determine significant differences between the means of the two time points. To see if there was a relationship between ΔH^+^ and carbon isotope, we plotted the mean species ΔH^+^ against the mean δ^13^C and ran a weighted least squares regression.

## RESULTS

### δ^13^C survey suggests the presence of CAM

To test for CAM across *Pinguicula*, we surveyed carbon isotope composition (δ^13^C) in 56 species spanning the genus, including 36 from the Mexican Clade. We also gathered previously published δ^13^C for *Pinguicula* and included those in our results presented in Table 1. Most *Pinguicula* species had a C_3_-like δ^13^C, ranging from -22 to -33‰ (Table 1). Notably, two species in the Mexican Clade, *P. agnata* (mean δ^13^C = -18.65‰ ± 0.58 *s.e.*, n = 4) and *P. kondoi* (mean δ^13^C = -17.62‰ ± 1.42 *s.e.*, n = 3) had mean δ^13^C less negative than -20‰ suggesting the presence of a carbon concentrating mechanism, that is, CAM. While the isotopic survey heavily implied the presence of CAM in these two Mexican *Pinguicula* species, they could not rule out the possibility that species in the C_3_ isotopic range demonstrate CAM intermediacy, and therefore those required further physiological validation (Winter et al., 2005; Messerschmid et al., 2021).

**Table 1.**
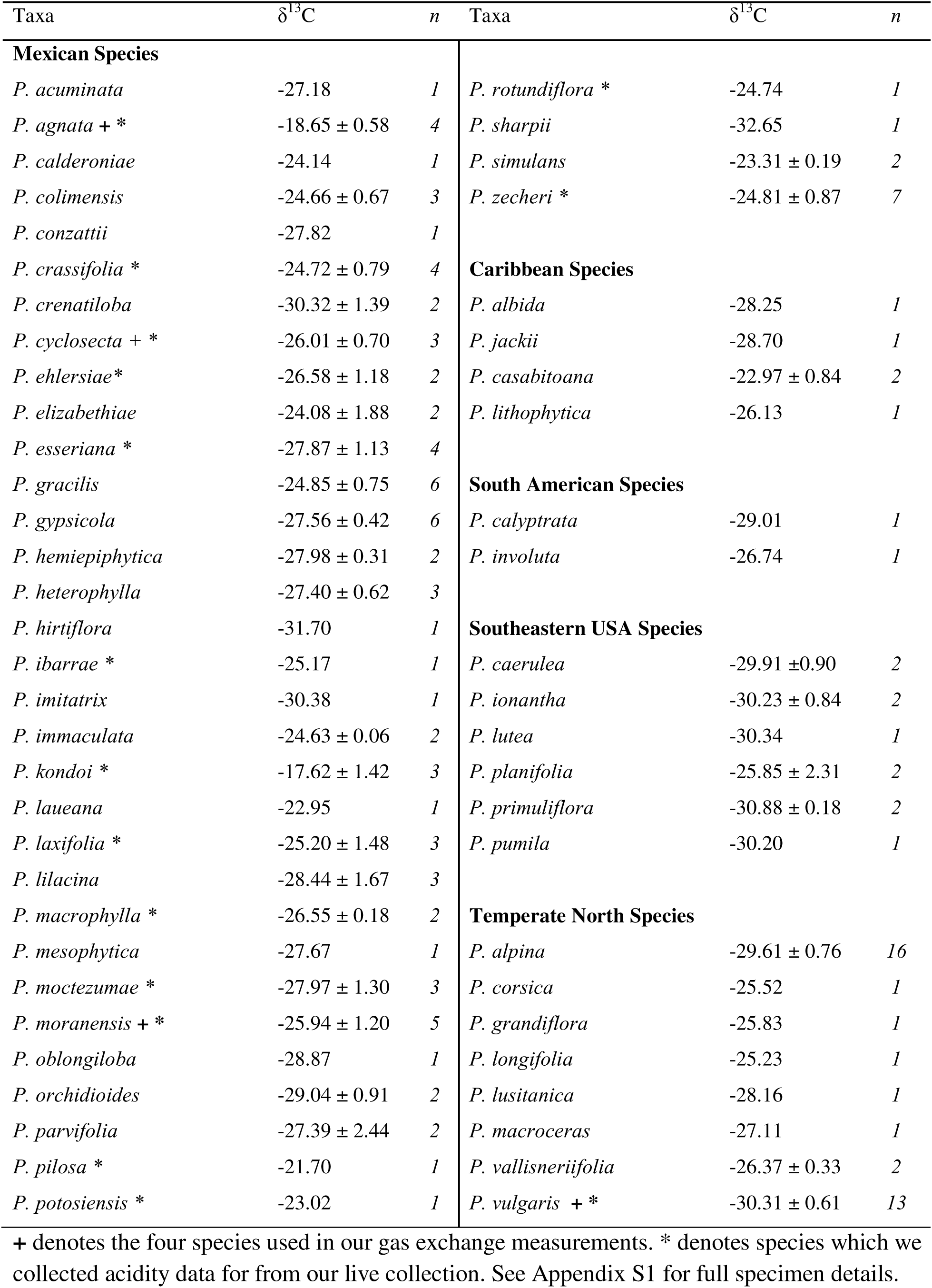
δ^13^C values from *Pinguicula* species with an emphasis on Mexican Species.

### Gas Exchange Measurements reveal diversity of CAM

To test CAM variation in *Pinguicula*, we selected representative species for our gas exchange measurements based on carbon isotope ratios that spanned C_3_-like to CAM-like values that therefore may reflect variation along the CAM continuum (Table 1). Based on the δ^13^C values (mean δ^13^C = -30.31‰ ± 0.61 s.e., n = 13), we did not expect to see evidence for CAM in *Pinguicula vulgaris*, our temperate species. This hypothesis was supported by the temporal pattern of net CO assimilation (*A*), which showed no increase during the dark period that would indicate nocturnal carbon uptake (Figure 2A). Additionally, net positive *A* was restricted to the light period, and our estimated proportion of CO_2_ recycled via CAM ranged from -0.23 to 2.48 % (Table 2).

**Figure 2:**
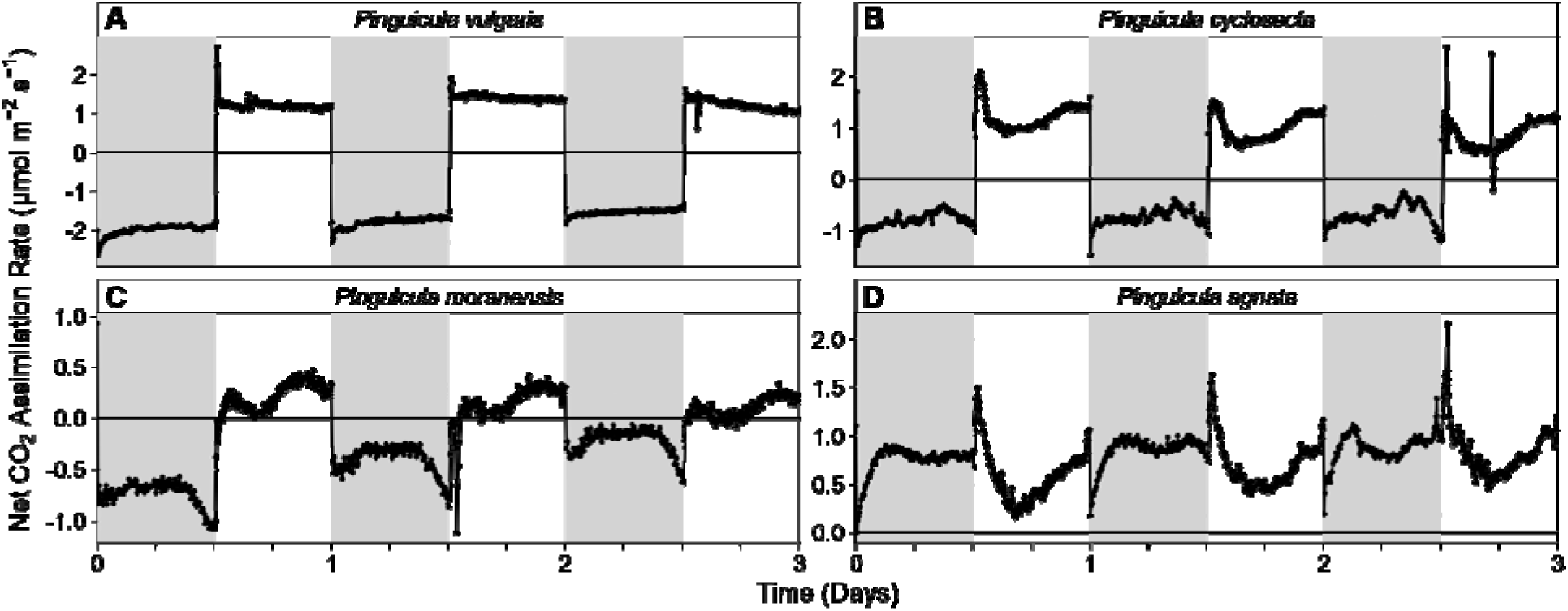
Net CO_2_ assimilation rate of four *Pinguicula* species measured continuously over three days. Grey shaded areas indicate dark periods; white shaded areas indicate light periods. Water was withheld starting Day 0.

**Table 2.** Estimates of 24 hr net CO_2_ assimilation in *P. agnata* and recycled respiratory CO_2_ in *P. cyclosecta*, *moranensis*, and vulgaris.

| Species | Day | Integrated Net CO<sub>2</sub> assimilation (mmol m<sup>-2</sup>) |  | Daily Net CO<sub>2</sub> assimilation (mmol m<sup>-2</sup>) | Proportion of net CO<sub>2</sub> acquired via Dark (%) |
| --- | --- | --- | --- | --- | --- |
|  |  | Dark (12 hr) | Light (12 hr) | Day (24 hr) |  |
| *P. agnata* | 1 | 31.77 | 25.56 | 57.33 | 55.52 |
| *P. agnata* | 2 | 36.60 | 30.49 | 67.09 | 54.55 |
| *P. agnata* | 3 | 37.44 | 36.15 | 73.59 | 50.87 |
| Species | Dark Period | Integrated Net CO<sub>2</sub> assimilation (mmol m<sup>-2</sup>) |  | Estimated CO<sub>2</sub> recycled via CAM (mmol m<sup>-2</sup>) | Proportion of respired CO<sub>2</sub> recycled via CAM (%) |
|  |  | Observed | Estimated |  |  |
| *P. cyclosecta* | 1 | -34.22 | -39.06 | 4.83 | 12.37 |
| *P. cyclosecta* | 2 | -31.44 | -36.02 | 4.58 | 12.72 |
| *P. cyclosecta* | 3 | -29.39 | -38.51 | 9.12 | 23.67 |
| *P. moranensis* | 1 | -31.46 | -37.64 | 6.18 | 16.42 |
| *P. moranensis* | 2 | -17.25 | -24.90 | 7.55 | 30.44 |
| *P. moranensis* | 3 | -9.37 | -16.57 | 7.21 | 43.49 |
| *P. vulgaris* | 1 | -85.21 | -87.12 | 1.90 | 2.18 |
| *P. vulgaris* | 2 | -76.63 | -78.58 | 1.95 | 2.48 |
| *P. vulgaris* | 3 | -65.41 | -65.26 | -0.15 | -0.23 |

729 All values in this table are extrapolated from the gas exchange data presented in Figure 2.

*Pinguicula cyclosecta* (mean δ^13^C = -26.01‰ ± 0.70 *s.e.*, n = 3) and *P. moranensis* (mean δ^13^C = -25.94‰ ± 1.20 *s.e.*, n = 5), both with δ^13^C in the C_3_ range, demonstrated evidence for CAM (Figure 2B, C). In both, we observed a non-constant, concave shape of *A* in the dark period, although never positive. In the dark period, the times of greatest net CO_2_ loss (i.e., most negative *A*) were observed at the beginnings and ends of the dark periods. The time of lowest net CO_2_ loss was observed in the middle of the dark period, consistent with CAM-like PEPc activity recycling respiratory CO_2_ (Griffiths et al., 1989). Over the three days of measurements, *P. moranensis* showed a decrease in dark net CO_2_ loss with each consecutive night, while the shape of the dark period *A* retained the characteristic CAM curvature (Figure 2C, Table 2). *P. cyclosecta* also showed a decrease in dark net CO_2_ loss, but between the second and third night (Figure 2B, Table 2). For both individuals, we observed a transient early light period decline in *A* before increasing later in the day suggesting stomatal closure and opening respectively. This occurred consistently with each beginning of the light period.

*Pinguicula agnata*, one of the species with a CAM-like δ^13^C (mean δ^13^C = -18.65‰ ± 0.58 *s.e.*, n = 4), showed evidence for Full CAM physiology (Figure 2D). *A* remained positive in both the light and dark periods across all three days of measurement, never falling below zero. *A* increased towards the middle of the night and declined slightly towards the end of the dark period, showing the characteristic concave shape of CAM. At the transition period between the dark and light period, there was a brief, but pronounced spike in *A* observed each day before rapidly declining to lower sustained rates. Assimilation then gradually increased again toward the end of the light period. Across the three measured days, dark period *A* consistently accounted for greater than 50% of total daily carbon gain (Table 2), demonstrating that CAM contributes a majority of carbon assimilation, even under well-watered conditions.

Continuous gas exchange measurements of *P. martinezii* over a 20-day drought time course revealed substantial changes in the diurnal pattern of *A* over the course of water-withholding and subsequent rewatering (Figure 3). At the start of the measurements, light period net CO_2_ assimilation rates were positive, while dark period rates were negative. Despite this, there was an observable curvature to the dark period *A*. Through days 1–5 of water-withholding, this general pattern was maintained. Starting on day 6 onward, light period *A* rapidly declined with each subsequent day. Conversely, dark period *A* became less negative with each day, approaching zero by approximately day 9. From day 10 onward, dark period *A* was net positive for the majority of the night, though maintaining the curvature and peaking in the middle of the dark period. Following rewatering on day 14, there was one day of strongly negative dark period *A* and near zero light period rates. However, from day 16 onwards, dark period *A* returned to the concave pattern resembling before water-withholding and light period rates gradually increased until near full recovery.

**Figure 3:**
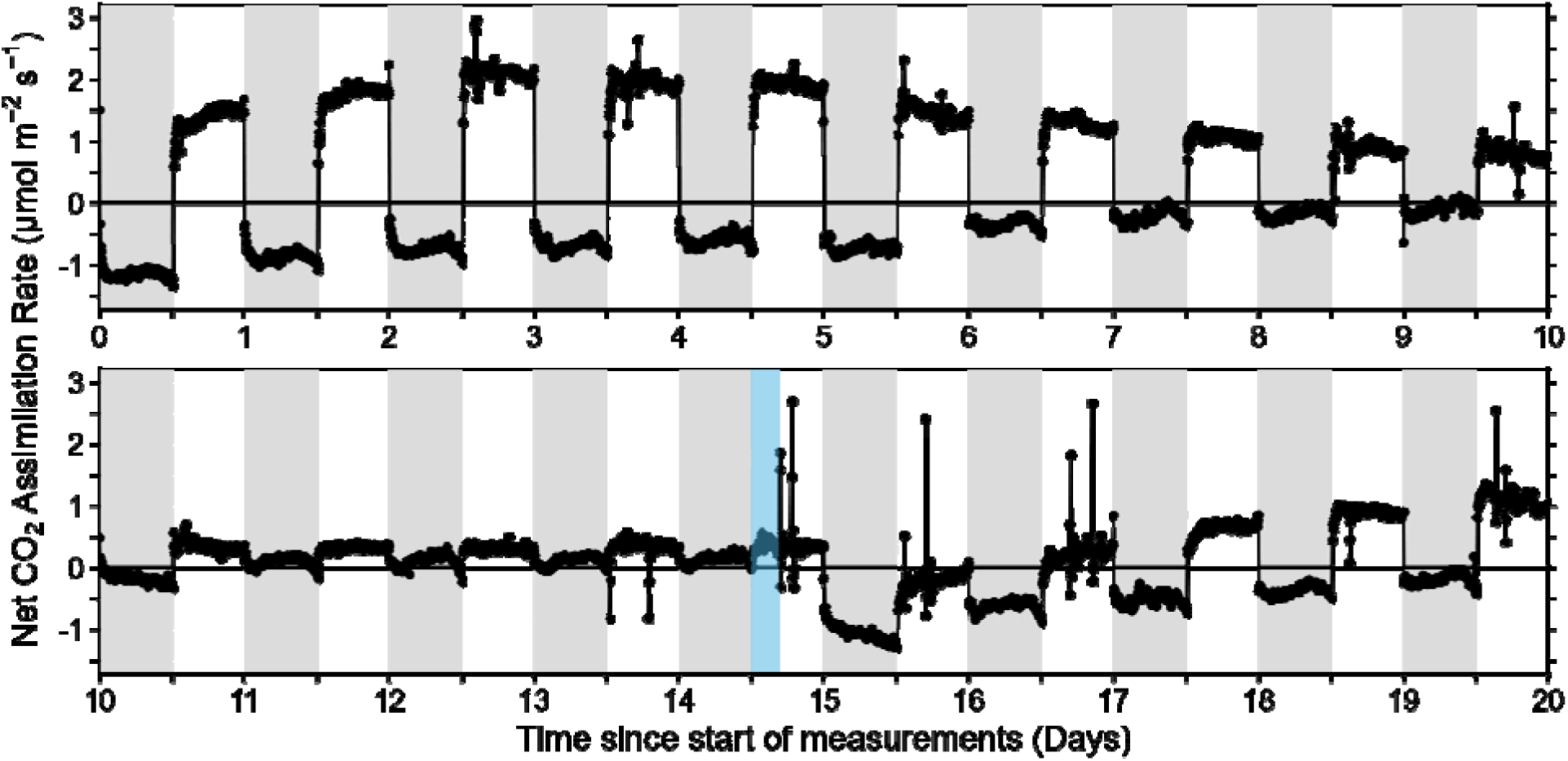
Net CO_2_ assimilation rate of *Pinguicula martinezii* measured continuously over a 20 day period shows facultative induction of CAM. Grey shaded areas indicate dark periods; white shaded areas indicate light periods. Water was withheld starting Day 0 and the plant was re-watered in the light period of the 14th day indicated by the blue shaded region.

Overall, our gas exchange experiments on living specimens supported variability in the patterns of carbon assimilation in *Pinguicula*, and the presence of a CAM continuum. We found evidence for C_3_, low levels of CAM, facultative CAM, and constitutive CAM.

### Nocturnal Acid Accumulation Varies Across *Pinguicula*

To increase the diversity of our sampling and complement our δ^13^C and gas exchange data, we investigated nocturnal acid accumulation in specimens from our living collections. Of the 20 species we sampled for overnight changes in tissue acidity, 16 showed significantly different means (Welch’s t-test, alpha = 0.05) between dawn and dusk (Figure 4). Among the species selected for gas exchange, the magnitude of this shift varied considerably. *P. agnata* exhibited the largest mean difference in titratable acidity (ΔH^+^ = 20.94 ± 0.90 standard error of difference (SED) µmol H^+^ g ¹ FW, t = 23.20, df = 4.00, P = 0.00002), roughly three times that of *P. cyclosecta* (ΔH^+^ = 7.11 ± 0.84 SED µmol H^+^ g ¹ FW, t = 8.39, df = 2.94, P = 0.0038) and *P. moranensis* (ΔH^+^ = 7.62 ± 0.98 SED µmol H^+^ g ¹ FW, t = 7.71, df = 2.55, P = 0.0078), which were broadly comparable to one another. *P. vulgaris*, by contrast, showed no significant overnight change in acidity (ΔH^+^ = 0.94 ± 1.03 SED µmol H^+^ g ¹ FW, t = 0.91, df = 2.98, P = 0.43). Of the remaining species, three had non-significant ΔH^+^: *P. macrophylla* (ΔH^+^ = 0.22 ± 0.76 SED µmol H^+^ g ¹ FW, t = 0.30, df = 3.94, P = 0.78), *P. moctezumae* (ΔH^+^ = 2.14 ± 0.97 SED µmol H^+^ g ¹ FW, t = 2.22, df = 3.13, P = 0.11), and *P. potosiensis* (ΔH^+^ = 2.42 ± 2.21 SED µmol H^+^ g ¹ FW, t = 1.10, df = 2.19, P = 0.38).

**Figure 4:**
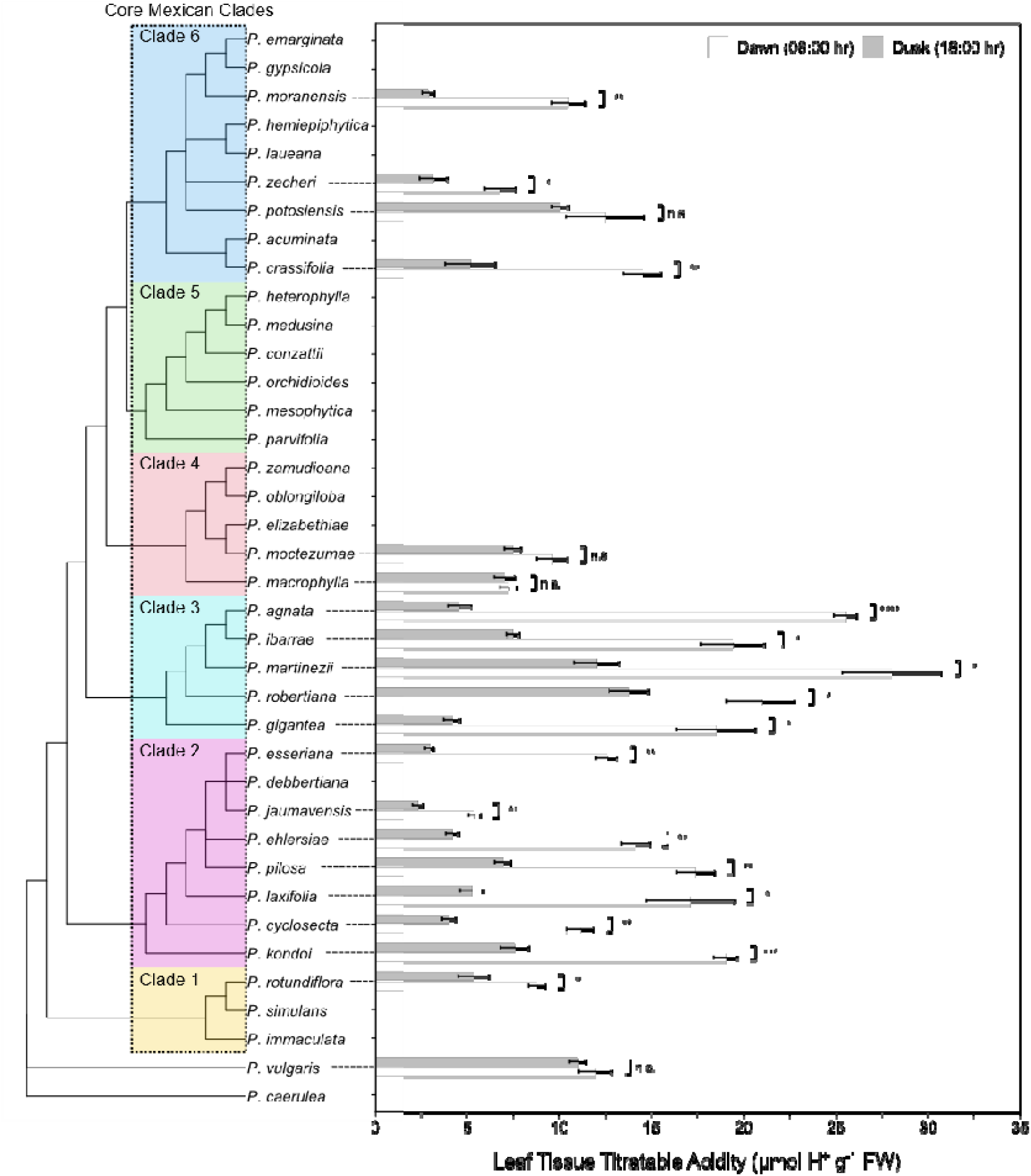
Leaf Titratable Acidity at Dusk and Dawn in 20 *Pinguicula* species and their phylogenetic placement. The core Mexican Clades are highlighted and numbered. Topology and clade numbering adapted from (Liu et al., 2026) with additional taxa appended as a polytomy from (Shimai et al., 2021). White and grey bars represent mean tissue acidity ± standard error (n = 3) at dawn (06:00 h) and dusk (18:00 h) respectively. For each species, significant differences between dawn and dusk were determined with a Welch’s t-test. Given the exploratory species-by-species nature of the figure, no correction for multiple comparisons was applied.

We then explored whether there were relationships between δ^13^C and ΔH^+^. Of the 20 species with available acidity data, 16 also had available δ^13^C data. Across these 16 examined species, the species mean δ^13^C was significantly and positively associated with the species mean ΔH^+^ (y = 1.07x + 34.51, R² = 0.38, P = 0.006; Figure 5). This indicated that species with less negative (i.e., more enriched) δ^13^C tend to exhibit larger diel shifts in tissue acidity, which is consistent with greater CAM expression.

**Figure 5:**
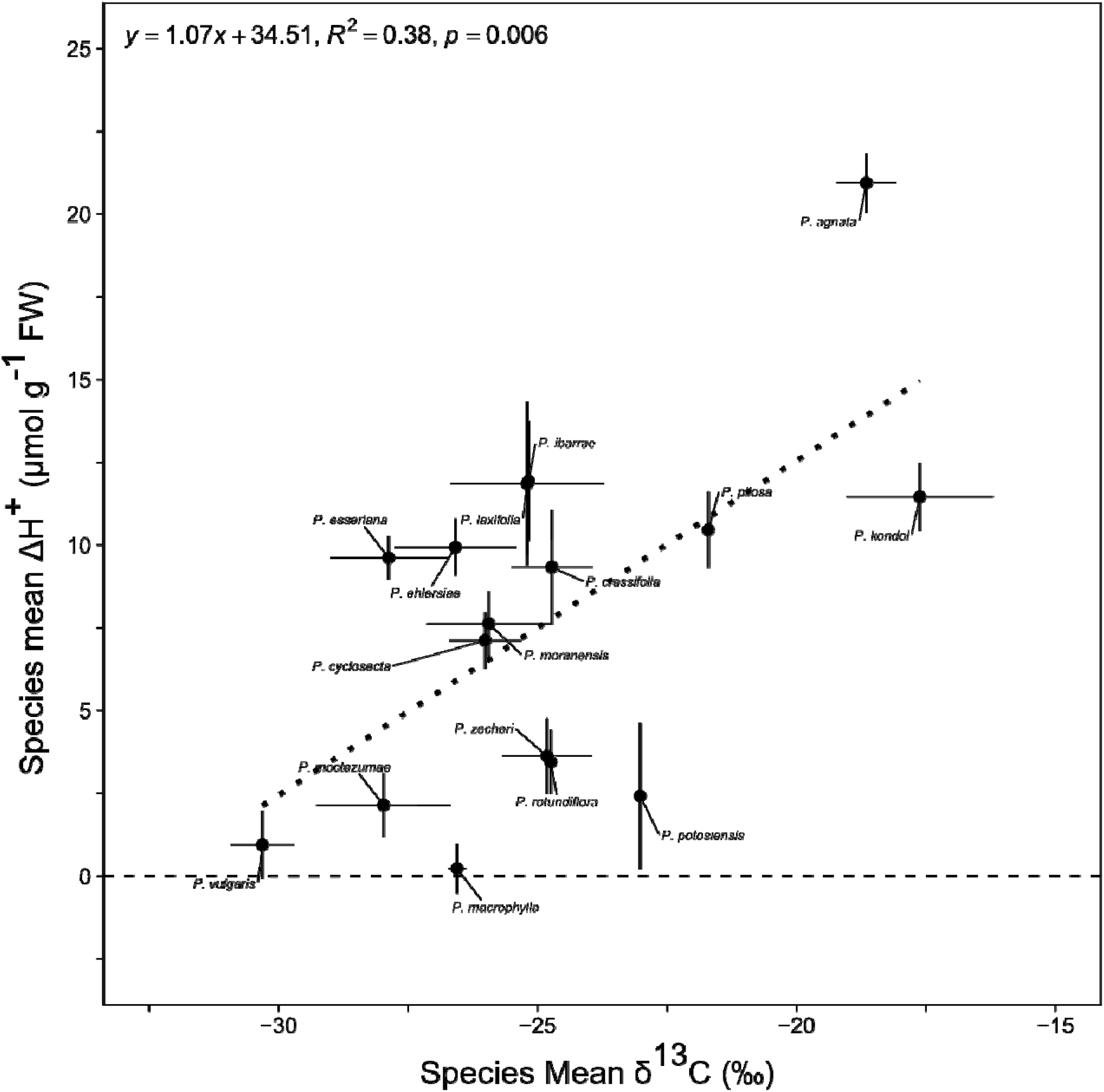
Relationship between species mean δ¹³C (‰) and species mean ΔHD (µmol HD gD¹ FM) across 16 *Pinguicula* species. Each point represents the species mean, with horizontal bars indicating standard error for δ¹³C and vertical bars representing the standard error of the difference in the mean of H. The dotted line shows the fitted weighted least squares regression (y = 1.07x + 34.51, R² = 0.38, P = 0.006), and the shaded grey region represents the 95% confidence interval around the regression line.

## DISCUSSION

Across our genus-wide survey of carbon isotopes, and measurements of gas exchange, and titratable acidity in our living collections, *Pinguicula* shows evidence of a CAM continuum. Our δ^13^C survey of 56 species identified two isotopically CAM-like species, while gas exchange revealed a range of physiologies. Titratable acidity measurements across 20 species largely corroborated these patterns, with 16 species showing significant day-night acidity shifts and a significant positive relationship between δ^13^C and ΔH^+^ across the tested taxa.

### Placing *Pinguicula* species on the CAM Continuum

Despite the fluid nature of the CAM continuum, three broadly defined groups can be used for comparative purposes: (1) non-CAM plants where the CAM pathway is never used in the plant’s lifespan, and the plant lacks the capacity to perform any amount of CAM, such as in fully C_3_ or C_4_ plants; (2) C_3_-CAM or CAM Intermediate, where low-levels of CAM constitutes a minority (< 50%) of the plant’s carbon economy (including recycling of respiratory CO_2_) and the rest comes from C_3_ (or C_4_) photosynthesis (C_3_+CAM in (Edwards, 2019), ‘minority CAM’ in (Gilman et al., 2024)); and (3) Strong CAM or ‘Full CAM’ plants, where CAM contributes a majority of the total net carbon economy (‘primary CAM’ in (Gilman et al., 2024), ‘CAM plant’ in (Winter, 2019)). Here we will use non-CAM, C_3_-CAM, and ‘Full CAM’ to delineate our groups.

Our study reveals that species in *Pinguicula* span the CAM continuum. Specifically, we find evidence for non-CAM, C_3_-CAM, and Full CAM species within the genus based on our measurements of net CO_2_ assimilation (*A*). *Pinguicula vulgaris,* our temperate representative, showed no evidence for CAM under well-watered conditions, consistent with a purely C_3_ physiology (Figure 2). This aligns with the δ^13^C values in the C_3_ range, lack of nocturnal tissue acidification, and inability to survive even moderate drought. We found that the *P. vulgaris* leaves senesce after more than 4 days of water-withholding, despite the more water-retentive potting mix. Therefore, on the CAM continuum, *P. vulgaris* is placed in the non-CAM extreme. Our results are also consistent with prior titratable acidity measurements in the closely related temperate species *P. grandiflora*, which showed no evidence for nocturnal acid accumulation (Fleck et al., 2026) together supporting C_3_ as the ancestral physiological state in *Pinguicula*.

Despite δ^13^C values in the C_3_ range, our individuals of the Mexican species *P. cyclosecta*, and *P. moranensis*, showed patterns of *A* inconsistent with purely C_3_ physiology (Figure 2B, C). The dark period curvature of *A* is a recognized signature of low amounts of CAM (Winter and Holtum, 2015), reflecting the recycling of respiratory CO_2_ at night rather than net carbon gain. This pattern has previously been observed in other C_3_-CAM species such as *Bulnesia*, *Pilea*, and *Jatropha* (Winter and Holtum, 2015; Winter et al., 2021; Mok et al., 2023). The transient early light period decline in *A* suggests partial stomatal closure, common in many CAM plants (Cushman, 2001; Males and Griffiths, 2017). This places both *P. cyclosecta* and *P. moranensis* within the C_3_-CAM range when viewed under the CAM continuum. While they constitutively use CAM, they operate at the lower end of CAM expression where CAM does not result in net carbon gain, consistent with the estimations of CAM recycled respiratory CO_2_ of 12.37 to 23.67% in *P. cyclosecta* and 16.42 to 43.49% in *P. moranensis* (Table 2). The progressive decrease in dark period CO_2_ loss seen across consecutive days in *P. moranensis*, but not in *P. cyclosecta*, suggests that *P. moranensis* also has a facultative component to its CAM expression. Given our measurements only lasted three days, it is not clear whether *P. moranensis*, if stressed long enough, could switch to positive *A* at night. Facultative CAM cannot be ruled out in *P. cyclosecta*, as it is possible that water was not withheld long enough to elicit a substantial response to drought. *P. agnata* demonstrated physiology consistent with Full CAM. *A* remained positive throughout both the light and dark periods, with dark period *A* accounting for a substantial share of total daily carbon gain (Figure 2D). The pronounced spike and dip in *A* at the dark-to-light period transition suggests stomatal closure. Once the nocturnally stored acids are depleted, stomata reopen later in the day to capture atmospheric CO_2_ before the dark (Males and Griffiths, 2017). These gas exchange patterns have previously been observed in several other CAM lineages (Lüttge, 2004). Combined with δ^13^C values above the -20‰ threshold, and the highest ΔH^+^ of all tested species, our combined evidence firmly places *P. agnata* on the Full CAM range of the CAM continuum. *P. agnata* challenges conventional associations of Full CAM with structurally robust succulents such as in cacti and many orchids (Gilman and Edwards, 2020).

The small stature and delicate succulent leaves of *P. agnata* suggest Full CAM may be more taxonomically and morphologically widespread than currently appreciated. We speculate this could partly reflect a bias towards screening CAM in plants with classically sturdy, robust morphology, potentially overlooking Full CAM in more structurally delicate taxa.

Previous reports of titratable acidity in *P. martinezii* support C_3_ physiology (Fleck et al., 2026) our gas exchange measurements and significant titratable acidity (ΔH^+^) under well-watered conditions suggest that it is CAM. Under well-watered conditions, patterns of *A* resembled other low-level C_3_-CAM plants, in which light period *A* remains positive as seen in typical C_3_ plants, and dark period *A* has the characteristic curvature (Winter, 2019; Winter et al., 2021; Mok et al., 2023). Water-withholding drove dark period *A* to gradually increase until exceeding zero, accompanied by declines in the light period *A* (Figure 3). Rewatering restored the well-watered pattern within two days. Since *P. martinezii* fixes little carbon via CAM under well-watered conditions and only becomes net positive after 14 days of water-withholding, we place it as a C_3_-CAM on the CAM continuum.

The majority of species we sampled for titratable acidity (16 of 20) showed significant changes in tissue acidity overnight, indicating some degree of nocturnal acid accumulation is present in many species across the Mexican Clade. Across 16 species for which both δ^13^C and ΔH^+^ were available, the positive relationship between species mean δ^13^C and species mean ΔH^+^ indicates that carbon isotope enrichment and diel organic acid fluctuation are coupled in Mexican *Pinguicula* (Figure 5). *P. vulgaris*, our C_3_ representative outgroup, anchors the low end of both axes with the most negative δ^13^C and minimal ΔH^+^. Both *P. cyclosecta* and *P. moranensis*, despite falling well within the C_3_ δ^13^C range, show modest ΔH^+^ consistent with our gas exchange evidence suggesting low amounts of CAM. At the upper limit, *P. agnata* shows substantially higher ΔH^+^ than predicted, consistent with its designation as a Full CAM plant.

These results provide additional support for a CAM continuum in *Pinguicula* and challenge the typical bimodal distribution of isotope ranges. *Pinguicula* suggests that if sampling was better in CAM-evolving lineages, there may be a multi-modal rather than bi-modal isotopic distribution.

### Insights into CAM Evolution from *Pinguicula*

Our identification of *Pinguicula* species spanning the variation of CAM provides a valuable framework for understanding CAM evolution within a single lineage. Two models have been proposed for the CAM evolutionary trajectory: (1) a continuous, gradual transition from C_3_ to CAM (Silvera et al., 2009; Bräutigam et al., 2017); and (2) a stepwise model with evolutionary thresholds between discrete phenotypes analogous to the staged evolution proposed for C_4_ photosynthesis (Edwards, 2019). Our data are consistent with the view that these two models are perhaps not mutually exclusive (Edwards, 2023) and support that the phenotypic space between C_3_ and Full CAM in *Pinguicula* is both a continuum of CAM expression, and also contains at least three recognizable, functionally distinct states.

Firstly, there are non-CAM plants such as *P. vulgaris*, where CAM is not detectable and plants rapidly senesce under water withholding. Next, we observed a key difference from non-CAM to C_3_-CAM, where C_3_ photosynthesis still makes up most of a plant’s CO_2_ assimilation, but CAM contributes a minority to the net-carbon economy, as in *P. cyclosecta* and *P. moranensis*.

This C_3_-CAM phenotype could provide increased growth through the recycling of respiratory CO_2_, reducing net carbon loss (Hartinger et al., 2026). Another variation of C_3_-CAM physiology was observed in *P. martinezii*, which constitutively recycles respiratory CO_2_ with CAM, even under well-watered conditions, and further upregulates CAM to a point of net positive CO_2_ assimilation under drought. Lastly, a core distinction between C_3_-CAM and Full CAM, is the obligatory reliance on CAM for net growth. This is seen in *P. agnata*, where even under well-watered conditions, CAM constitutively accounts for over 50% of total CO_2_ assimilation.

We propose three forces that could be associated with the evolution of CAM in *Pinguicula*. Firstly, since carnivorous plants tend to have high respiration rates (Adamec, 2010, 2013; Pavlovič et al., 2022), they may operate close to a carbon-balance deficit, particularly under low-light or nutrient-poor conditions where leaf construction costs are high relative to the photosynthetic return (Ellison and Adamec, 2011). Under such conditions, constitutive C_3_-CAM as seen in *P. cyclosecta* and *P. moranensis*, could meaningfully offset this deficit by recycling respiratory CO_2_ that would otherwise be lost at night. Although the amount of carbon recovered each night may be modest, the cumulative respiratory CO_2_ recycled over the lifetime of the leaf could substantially outweigh the metabolic costs of maintaining constitutive low-level CAM. Secondly, this carbon-conservation benefit could be enhanced in the seasonally dry habitats occupied by Mexican *Pinguicula*, where daytime stomatal closure due to limited water loss would severely restrict carbon gain. In such contexts, nocturnal CO_2_ assimilation via PEPc not only conserves respired CO_2_ but allows continued carbon assimilation while stomata are closed during the heat of the day. The dual advantage of respiratory carbon recycling and increased water-use-efficiency could be the reason for the transition to Full CAM and relying on nocturnal CO_2_ fixation for net growth. Lastly, a third benefit could be photoprotection. Since CAM plants fix CO_2_ at night and refix it during the day, the Calvin cycle can continue operating as an electron sink even when stomata close in response to heat or water stress. This limits the excess excitation energy that would otherwise drive photoinhibition (Ceusters et al., 2019). The capacity to sustain some carbon-linked electron flow behind closed stomata has been shown to protect the photosynthetic apparatus during drought in *Pereskia aculeata* (de Barros et al., 2024). For Mexican Pinguicula, which have delicate leaves and exposure to high irradiance on open rock faces and cliffs, this photoprotective function of CAM could be just as important as its water-conservation or carbon-recycling benefits. We additionally hypothesize that the evolution of CAM could have enabled the adoption of this lithophytic or epiphytic lifestyle in Mexican *Pinguicula*. Rock crevices and cliff faces are characterized by minimal substrate volume, rapid drainage, and low water-holding capacity: all conditions that intensify water stress. The shift to growing in such habitats may have placed Mexican lineages under sustained selection for traits that improve water-use-efficiency, with Full CAM being one possible outcome.

The temperate *Pinguicula* species, which are early-diverging within the genus (Shimai et al., 2021), inhabit wetlands and bogs with abundant year-round water. Since the measured physiology of *P. vulgaris* is consistent with C_3_ photosynthesis, this supports C_3_ as being ancestral to the genus, a conclusion also reinforced by prior findings of a lack of nocturnal acid accumulation in *P. grandiflora* (Fleck et al., 2026). Our results suggest CAM arose in the Mexican Clade, thus resolving the origin of CAM within the Mexican Clade should be a future priority that will require targeted sampling of key taxa, such as *P. lilacina* and *P. sharpii*, based on their early-diverging phylogenetic placement and divergent ecology. These annual taxa from water-rich habitats are sister to all other Mexican species (Shimai et al., 2021), and will thus be particularly informative. If these species lack CAM, it would shift the origin of CAM more recently into the core Mexican Clades and strengthen the case for a more recent, ecologically driven transition. Conversely, evidence of even weak CAM in these taxa could imply an earlier origin and a longer trajectory of phenotypic elaboration within the clade, in which case, CAM could be considered an exaptation preceding expansion into water-limited niches. Along with these important taxa, more thorough sampling and physiological surveying of clades 4 and 5 (Figure 4) will also be required. The lack of significant nocturnal acid accumulation in our two species from Clade 4 suggests the possibility of a reversion to C_3_ physiology. Species in Clade 4 are found in more mesic, humid habitats (Lampard et al., 2016) and so additional testing involving water withholding should be done to investigate the presence of facultative CAM. This would help clarify whether there is a reversion to C_3_ or multiple evolutionary origins of CAM within the Mexican Clade.

The diversity of CAM phenotypes we observed in this genus of small, striking carnivorous plants is representative of the immense variation of CAM found across the land plants. This is reminiscent of patterns seen in other lineages where CAM has been linked to ecological diversification and radiation, including in the Bromeliaceae (Crayn et al., 2015) and Orchidaceae (Givnish et al., 2015), where CAM has been associated with the occupation of epiphytic niches and water is a limiting resource. Moreover, our observations from this study raise the possibility that the evolution of CAM, and the subsequent diversification of its expression along the CAM continuum, contributed to the species radiation of Mexican *Pinguicula* by opening up a broader range of water-limited microhabitats to the clade.

## CONCLUSIONS

In our study, by combining carbon isotope surveys, gas exchange measurements, and titratable acidity across dozens of *Pinguicula* species, we demonstrate that *Pinguicula* spans the CAM continuum within a single genus. This includes non-CAM in our temperate outgroup species, to a variety of C_3_-CAM (both constitutive and facultative) and Full CAM in the Mexican Clade. Our results suggest that C_3_ is ancestral to *Pinguicula* and CAM evolved in the Mexican Clade. We propose the initial transition likely began as a constitutive, low-level mechanism for photoprotection or recycling respiratory CO_2_ in seasonally water-limited environments, which was elaborated into net growth via CAM in more xeric, lithophytic habitats. Future work to investigate the evolutionary trajectory of CAM should include increased physiological sampling of early-diverging Mexican taxa, such as *P. lilacina* and *P. sharpii*, along with following up with isotopically CAM-like species such as *P. kondoi*. Additionally, our data, together with the proposal that CAM offsets the elevated respiratory costs characteristic of carnivorous plants (Adamec, 2010, 2013; Ellison and Adamec, 2011), suggest that the existing cost-benefit frameworks for plant carnivory, which typically weigh nutrient capture against photosynthetic carbon gain, could benefit from treating photosynthetic pathway itself as a further axis of variation. Given our hypothesis on the benefits of recycling respiratory CO_2_ in carnivorous plants, it may also be worth surveying other carnivorous plant lineages for evidence of low-level CAM expression.

Beyond resolving where and when CAM arose, the continuum described here opens several broader avenues for future work in *Pinguicula.* Because these physiologically distinct states occur among relatively closely related species in a monophyletic lineage, comparative genomics and transcriptomics between *Pinguicula* species with differing CAM physiologies could identify regulatory changes that accompany the transition from C_3_ to C_3_-CAM to Full CAM. Extending this comparative approach to the many other independent origins of CAM across land plants could also test whether such transitions rely on parallel molecular solutions or distinct, lineage-specific routes to a convergent physiological outcome.

## ACKNOWLEDGEMENTS

We thank Ian S. Gilman for thoughtful discussions throughout the development and progress of this research. We would like to thank Rainbow Carnivorous Plants, Sarracenia Northwest, Boggystyle, and Jason Hupp for supplying plants for these experiments. We also thank Cody Keilen and Nick Deason, for managing the growth chamber facilities at MSU.

## AUTHOR CONTRIBUTIONS

DM, KJG, and RV conceived of the project. DM conducted the experiments, did data analysis, and wrote the initial draft for the manuscript. KJG and RV provided comments and revised the manuscript. This work was partially funded by the Botanical Society of America Graduate Student Research Award and a Michigan State University Plant Resilience Institute Seed Grant to DM, as well as startup funds from Michigan State University to KJG.

## DATA AVAILABILITY

All the data that supports the findings of this study are available in the supplementary materials of this article.

## Notes

### Competing Interest Statement

The authors have declared no competing interest.

## LITERATURE CITED

Adamec, L. 2013. A comparison of photosynthetic and respiration rates in six aquatic carnivorous Utricularia species differing in morphology. Aquatic Botany 111: 89–94.

Adamec, L. 2010. Dark respiration of leaves and traps of terrestrial carnivorous plants: are there greater energetic costs in traps? Open Life Sciences 5: 121–124.

Adamec, L., I. Matušíková, and A. Pavlovič. 2021. Recent ecophysiological, biochemical and evolutional insights into plant carnivory. Annals of Botany 128: 241–259.

Arakaki, M., P.-A. Christin, R. Nyffeler, A. Lendel, U. Eggli, R. M. Ogburn, E. Spriggs, et al. 2011. Contemporaneous and recent radiations of the world’s major succulent plant lineages. Proceedings of the National Academy of Sciences of the United States of America 108: 8379–8384.

de Barros, J. P. A., M. C. Lima Neto, N. D. da Silva Brito, P. J. Herminio, H. R. B. Santos, A. do N. Simões, V. G. Nunes, et al. 2024. The C3-CAM shift is crucial to the maintenance of the photosynthetic apparatus integrity in Pereskia aculeata under prolonged and severe drought. Acta physiologiae plantarum 46.

Borland, A. M., V. A. Barrera Zambrano, J. Ceusters, and K. Shorrock. 2011. The photosynthetic plasticity of crassulacean acid metabolism: an evolutionary innovation for sustainable productivity in a changing world. The New Phytologist 191: 619–633.

Bräutigam, A., U. Schlüter, M. Eisenhut, and U. Gowik. 2017. On the evolutionary origin of CAM photosynthesis. Plant Physiology 174: 473–477.

Ceusters, N., R. Valcke, M. Frans, J. E. Claes, W. Van den Ende, and J. Ceusters. 2019. Performance index and PSII connectivity under drought and contrasting light regimes in the CAM orchid Phalaenopsis. Frontiers in Plant Science 10: 1012.

Crayn, D. M., K. Winter, K. Schulte, and J. A. C. Smith. 2015. Photosynthetic pathways in Bromeliaceae: phylogenetic and ecological significance of CAM and C3 based on carbon isotope ratios for 1893 species. Botanical Journal of the Linnean Society. Linnean Society of London 178: 169–221.

Crayn, D. M., K. Winter, and J. A. C. Smith. 2004. Multiple origins of crassulacean acid metabolism and the epiphytic habit in the Neotropical family Bromeliaceae. Proceedings of the National Academy of Sciences of the United States of America 101: 3703–3708.

Cushman, J. C. 2001. Crassulacean acid metabolism. A plastic photosynthetic adaptation to arid environments. Plant Physiology 127: 1439–1448.

Domínguez, Y., P. Temple, I. Pančo, and V. F. O. Miranda. 2024. Biogeographical patterns of Pinguicula L. (Lentibulariaceae) in the Americas revealed by endemicity and habitat suitability analyses. Flora 313: Not Available.

Edwards, E. J. 2019. Evolutionary trajectories, accessibility and other metaphors: the case of C4 and CAM photosynthesis. The New Phytologist 223: 1742–1755.

Edwards, E. J. 2023. Reconciling continuous and discrete models of C4 and CAM evolution. Annals of Botany 132: 717–725.

Ellison, A. M. 2006. Nutrient limitation and stoichiometry of carnivorous plants. Plant Biology (Stuttgart, Germany) 8: 740–747.

Ellison, A. M., and L. Adamec. 2011. Ecophysiological traits of terrestrial and aquatic carnivorous plants: are the costs and benefits the same? Oikos (Copenhagen, Denmark) 120: 1721–1731.

Ellison, A. M., and N. J. Gotelli. 2001. Evolutionary ecology of carnivorous plants. Trends in Ecology & Evolution 16: 623–629.

Farquhar, G. D., and J. Lloyd. 1993. Carbon and oxygen isotope effects in the exchange of carbon dioxide between terrestrial plants and the atmosphere. Stable Isotopes and Plant Carbon-water Relations, 47–70. Elsevier.

Fleck, N. J., T. F. E. Messerschmid, A. Fleischmann, R. C. Ferrari, and G. Kadereit. 2026. Yes, we CAM! First evidence of CAM photosynthesis in a carnivorous plant. Plant Biology (Stuttgart, Germany) 28: 272–281.

Gilman, I. S., and E. J. Edwards. 2020. Crassulacean acid metabolism. Current Biology 30: R57–R62.

Gilman, I. S., K. Heyduk, C. Maya-Lastra, L. P. Hancock, and E. J. Edwards. 2024. Predicting photosynthetic pathway from anatomy using machine learning. The New Phytologist 242: 1029–1042.

Gilman, I. S., J. A. C. Smith, J. A. M. Holtum, R. F. Sage, K. Silvera, K. Winter, and E. J. Edwards. 2023. The CAM lineages of planet Earth. Annals of Botany 132: 627–654.

Givnish, T. J. 2015. New evidence on the origin of carnivorous plants. Proceedings of the National Academy of Sciences of the United States of America 112: 10–11.

Givnish, T. J., E. L. Burkhardt, R. E. Happel, and J. D. Weintraub. 1984. Carnivory in the bromeliad Brocchinia reducta, with a cost/benefit model for the general restriction of carnivorous plants to sunny, moist, nutrient-poor habitats. The American Naturalist 124: 479–497.

Givnish, T. J., D. Spalink, M. Ames, S. P. Lyon, S. J. Hunter, A. Zuluaga, W. J. D. Iles, et al. 2015. Orchid phylogenomics and multiple drivers of their extraordinary diversification. *Proceedings*. Biological Sciences 282: 20151553.

Griffiths, H., B. L. Ong, P. N. Avadhani, and C. J. Goh. 1989. Recycling of respiratory CO2 during Crassulacean acid metabolism: alleviation of photoinhibition in Pyrrosia piloselloides. Planta 179: 115–122.

Hájek, T., and L. Adamec. 2010. Photosynthesis and dark respiration of leaves of terrestrial carnivorous plants. Biologia 65: 69–74.

Hartinger, C., E. N. Smith, and L. J. Sweetlove. 2026. Metabolic design considerations for recycling of respiratory CO2 in leaves. The Plant Journal: For Cell and Molecular Biology 127: e71003.

Heslop-Harrison, Y. 2004. Pinguicula L. The Journal of Ecology 92: 1071–1118.

Heyduk, K. 2022. Evolution of Crassulacean acid metabolism in response to the environment: past, present, and future. Plant Physiology 190: 19–30.

Heyduk, K., J. J. Moreno-Villena, I. S. Gilman, P.-A. Christin, and E. J. Edwards. 2019. The genetics of convergent evolution: insights from plant photosynthesis. Nature Reviews. Genetics 20: 485–493.

Horn, J. W., B. W. van Ee, J. J. Morawetz, R. Riina, V. W. Steinmann, P. E. Berry, and K. J. Wurdack. 2012. Phylogenetics and the evolution of major structural characters in the giant genus Euphorbia L. (Euphorbiaceae). Molecular Phylogenetics and Evolution 63: 305–326.

Klink, S., P. Giesemann, and G. Gebauer. 2019. Picky carnivorous plants? Investigating preferences for preys’ trophic levels - a stable isotope natural abundance approach with two terrestrial and two aquatic Lentibulariaceae tested in Central Europe. Annals of Botany 123: 1167–1177.

Lampard, S., O. Gluch, A. Robinson, A. Fleischmann, P. Temple, S. McPherson, A. Roccia, et al. 2016. Pinguicula of Latin America: 2. Redfern Natural History Productions, Poole, England.

Legendre, L. 2000. The genus*Pinguicula*L. (*Lentibulariaceae*): an overview. Acta botanica Gallica: bulletin de la Societe botanique de France 147: 77–95.

Liu, Y., Q. Lin, S. J. Fleck, M. Mata-Rosas, E. Ibarra-Laclette, and T. Renner. 2026. Phylogenomics reveals the evolution of floral traits associated with pollinators and pollinator-prey conflict within the carnivorous Pinguicula subgenus Temnoceras. American Journal of Botany 113: e70156.

Lüttge, U. 1983. Ecophysiology of Carnivorous Plants. Physiological Plant Ecology III, 489–517. Springer Berlin Heidelberg, Berlin, Heidelberg.

Lüttge, U. 2004. Ecophysiology of crassulacean Acid Metabolism (CAM). Annals of Botany 93: 629–652.

Males, J., and H. Griffiths. 2017. Stomatal biology of CAM plants. Plant Physiology 174: 550–560.

Mata-Rosas, M., J. Hernández-Rendón, and M. M. Salinas-Rodríguez. 2020. In pursuit of Mexican Pinguicula: a journey to northwestern Oaxaca. Carnivorous Plant Newsletter 49: 17–27.

Messerschmid, T. F. E., J. Wehling, N. Bobon, A. Kahmen, C. Klak, J. A. Los, D. B. Nelson, et al. 2021. Carbon isotope composition of plant photosynthetic tissues reflects a Crassulacean Acid Metabolism (CAM) continuum in the majority of CAM lineages. Perspectives in Plant Ecology, Evolution and Systematics 51: 125619.

Miloslav Studnička. 1991. Interesting succulent features in the Pinguicula species from the Mexican evolutionary centre. Folia geobotanica et phytotaxonomica 26: 459–462.

Mok, D., A. Leung, P. Searles, T. L. Sage, and R. F. Sage. 2023. CAM photosynthesis in Bulnesia retama (Zygophyllaceae), a non-succulent desert shrub from South America. Annals of Botany 132: 655–670.

Okita, T. W. 1996. Crassulacean acid metabolism: Biochemistry, ecophysiology and evolution. Plant Science: An International Journal of Experimental Plant Biology 120: 116.

Osmond, C. B. 1978. Crassulacean acid metabolism: A curiosity in context. Annual Review of Plant Physiology 29: 379–414.

Osmond, C. B., W. G. Allaway, B. G. Sutton, J. H. Troughton, O. Queiroz, U. Lüttge, and K. Winter. 1973. Carbon isotope discrimination in photosynthesis of CAM plants. Nature 246: 41–42.

Osmond, C. B., H. Ziegler, W. Stichler, and P. Trimborn. 1975. Carbon isotope discrimination in alpine succulent plants supposed to be capable of crassulacean acid metabolism (CAM). Oecologia 18: 209–217.

Pavlovič, A. 2022. Photosynthesis in carnivorous plants: From genes to gas exchange of green hunters. Critical Reviews in Plant Sciences: 1–16.

Pavlovič, A., J. Jakšová, M. Hrivňacký, and L. Adamec. 2022. Alternative or cytochrome? Respiratory pathways in traps of aquatic carnivorous bladderwort Utricularia reflexa. Plant Signaling & Behavior 17: 2134967.

Pavlovič, A., and M. Saganová. 2015. A novel insight into the cost-benefit model for the evolution of botanical carnivory. Annals of Botany 115: 1075–1092.

Roccia, A., O. Gluch, S. Lampard, A. Robinson, A. Fleischmann, S. McPherson, L. Legendre, et al. 2016. Pinguicula of the temperate north: 1. Redfern Natural History Productions, Poole, England.

Rueda-Almazán, J. E., V. M. Hernández, J. R. Alcalá-Martínez, A. Fernández-Duque, M. Ruiz-Aguilar, and R. E. Alcalá. 2021. Spatial and temporal differences in the community structure of endophytic fungi in the carnivorous plant Pinguicula moranensis (Lentibulariaceae). Fungal Ecology 53: 101087.

Sage, R. F., P.-A. Christin, and E. J. Edwards. 2011. The C(4) plant lineages of planet Earth. Journal of Experimental Botany 62: 3155–3169.

Schneider, C. A., W. S. Rasband, and K. W. Eliceiri. 2012. NIH Image to ImageJ: 25 years of image analysis. Nature Methods 9: 671–675.

Shimai, H. 2017. Taxonomy and conservation ecology of the genus Pinguicula L. (Lentibulariaceae). University of Kent.

Shimai, H., H. Setoguchi, D. L. Roberts, and M. Sun. 2021. Biogeographical patterns and speciation of the genus Pinguicula (Lentibulariaceae) inferred by phylogenetic analyses. PloS One 16: e0252581.

Silvera, K., K. M. Neubig, W. M. Whitten, N. H. Williams, K. Winter, and J. C. Cushman. 2010. Evolution along the crassulacean acid metabolism continuum. Functional Plant Biology 37: 995–1010.

Silvera, K., L. S. Santiago, J. C. Cushman, and K. Winter. 2009. Crassulacean acid metabolism and epiphytism linked to adaptive radiations in the Orchidaceae. Plant Physiology 149: 1838–1847.

Silvestro, D., G. Zizka, and K. Schulte. 2014. Disentangling the effects of key innovations on the diversification of Bromelioideae (bromeliaceae): Key innovations in bromelioideae. Evolution; International Journal of Organic Evolution 68: 163–175.

Smith, J. A. C., and K. Winter. 1996. Taxonomic distribution of crassulacean acid metabolism. Crassulacean Acid Metabolism, Ecological Studies: Analysis and Synthesis. Berlin, Heidelberg, New York NY, 427–436. Springer Berlin Heidelberg, Berlin, Heidelberg.

Winter, K. 2019. Ecophysiology of constitutive and facultative CAM photosynthesis. Journal of Experimental Botany 70: 6495–6508.

Winter, K., J. Aranda, and J. A. M. Holtum. 2005. Carbon isotope composition and water-use efficiency in plants with crassulacean acid metabolism. Functional Plant Biology 32: 381–388.

Winter, K., M. Garcia, A. Virgo, and J. A. C. Smith. 2021. Low-level CAM photosynthesis in a succulent-leaved member of the Urticaceae, Pilea peperomioides. Functional Plant Biology 48: 683–690.

Winter, K., and J. A. M. Holtum. 2015. Cryptic crassulacean acid metabolism (CAM) in Jatropha curcas. Functional Plant Biology 42: 711–717.

Winter, K., and J. A. M. Holtum. 2002. How closely do the delta(13)C values of Crassulacean Acid metabolism plants reflect the proportion of CO(2) fixed during day and night? Plant Physiology 129: 1843–1851.

Winter, K., and J. A. C. Smith. 2022. CAM photosynthesis: the acid test. The New Phytologist 233: 599–609.

Yang, X., J. C. Cushman, A. M. Borland, E. J. Edwards, S. D. Wullschleger, G. A. Tuskan, N. A. Owen, et al. 2015. A roadmap for research on crassulacean acid metabolism (CAM) to enhance sustainable food and bioenergy production in a hotter, drier world. The New Phytologist 207: 491–504.

Yuan, G., M. M. Hassan, D. Liu, S. D. Lim, W. C. Yim, J. C. Cushman, K. Markel, et al. 2020. Biosystems design to accelerate C3-to-CAM progression. Biodesign Research 2020: 3686791.

